# TopoStats – a program for automated tracing of biomolecules from AFM images

**DOI:** 10.1101/2020.09.23.309609

**Authors:** Joseph G. Beton, Robert Moorehead, Luzie Helfmann, Robert Gray, Bart W. Hoogenboom, Agnel Praveen Joseph, Maya Topf, Alice L. B. Pyne

**Author notes:** Corresponding author: Alice L. B. Pyne.

## Abstract

We present TopoStats, a Python toolkit for automated editing and analysis of Atomic Force Microscopy images. The program automates identification and tracing of individual molecules in circular and linear conformations without user input. TopoStats was able to identify and trace a range of molecules within AFM images, finding, on average, 90% of all individual molecules and molecular assemblies within a wide field of view, and without the need for prior processing. DNA minicircles of varying size, DNA origami rings and pore forming proteins were identified and accurately traced with contour lengths of traces typically within 10 nm of the predicted contour length. TopoStats was also able to reliably identify and trace linear and enclosed circular molecules within a mixed population. The program is freely available via GitHub (https://github.com/afm-spm/TopoStats) and is intended to be modified and adapted for use if required.

## 1. Introduction

The use of Atomic Force Microscopy (AFM) in structural biology has been increasing over the past 30 years; AFM is now a versatile and accessible technique for directly imaging single biomolecules. This has led to its adoption by a wide community of biomaterials scientists, biophysicists and structural biologists. The advances in the field were facilitated in large part by hardware development: A 100X increase in image acquisition times has allowed the visualisation of dynamic biological processes [1][2]. This has been coupled with the development of more sensitive imaging modes and probes that can resolve the double-helix of DNA [3], or the subunits of a macromolecular protein complex [4], using commercially available equipment. These complement what is perhaps the defining feature of the AFM, unique among other structural tools operating at sub-nanometre resolution (cryo-EM, X-ray crystallography): its capacity for imaging in liquid at physiological temperatures, where imaged (bio)molecules are active and free to explore their native conformational space, with the caveats that molecules need to be adsorbed on a solid substrate and that the AFM probe exerts a small force (often ∼0.1 nN) on the sample. The technique has facilitated studies in which biological processes are watched as “molecular movies”: examples of which include the observation of myosin walking along an actin filament [5], observing the structural changes in bacteriorhodopsin upon light exposure [6] and visualising the assembly of centromeres [1]. In addition to seeing these changes in molecular structure, direct imaging with the AFM facilitates the observation of rare molecular states and conformations within a snapshot of a heterogeneous population, for example visualising deviations in the DNA double-helix induced by supercoiling [7]. These unique features of the AFM make it a versatile structural biology tool that can operate either standalone and/or complementing other techniques such as cryo EM and X-ray crystallography, where rare conformations of molecules are obscured by averaging.

However, bio-AFM has arguably suffered from a lack of the kind community-led investment in image processing and analytical capability seen for other techniques, most recently in the cryo EM “resolution revolution” [8][9]. Contrary to cryo EM or X-ray Crystallography, there are relatively few free and open source softwares available for automated analysis, despite the importance of automated analysis for minimising selection bias and facilitating statistical analysis. This puts a restraint on the use of AFM as a quantitative imaging technique. When image processing tools are used in AFM studies, analysis is commonly manually repeated for each individual molecule within images. Tools that facilitate this include the Bruker Nanoscope analysis, ImageJ [10] or the open source AFM imaging software, Gwyddion [11]. Automation with these software is possible but can be restricted to image correction (Nanoscope) or require writing home-made scripts (Gwyddion and ImageJ). An additional complication is the variable quality of AFM images, which can significantly impact image analysis, as molecules that are aggregated, in close proximity, or poorly resolved may be difficult to separate and have their conformation partly obscured. This lack of available software, combined with the specific problems with AFM sample preparation, is highlighted by a number of AFM studies which have required development of home-built image processing softwares, often developed simultaneously by separate labs to address practically the same samples and problems [7][12][13].

To directly address these issues, and to nucleate a virtual area of shared analytical infrastructure within the bioAFM community, we have developed TopoStats - an open-source Python utility that combines AFM image correction, molecule identification and tracing into a single automated protocol. We use a Python implementation of Gwyddion (pygwy) [11] for rapid image correction, which we feed directly into our own Python modules for automated tracing and analysis of biomolecules, described here step-by-step. We use multiple DNA minicircle samples to demonstrate TopoStats is a reliable and accurate tool for automated single molecule identification and tracing, before demonstrating its versatility when applied to biological and biomimetic pores. We encourage the community to contribute to these tools (available at Github https://github.com/afmstats/TopoStats), and hope that this can be a starting step to link AFM image analysis to the growing tools freely available through Python distributions.

## 2. Methods

### 2.1 TopoStats Automated Image Analysis

#### 2.1.1 Purpose of TopoStats

TopoStats was developed to be a simple, easy to use and open-source program intended to function as both a fully operational pipeline for generalised AFM image processing and molecular tracing as well as a platform for the development of more complex and specialised image processing routines. TopoStats is implemented in Python 2.7 and makes use of the freely available Gwyddion [11], NumPy [14] and SciPy [15] Python libraries. Using Gwyddion functions, TopoStats supports all commercially used file formats making its use unrestricted for labs running commercial, and most homebuilt AFMs. We actively encourage and welcome community development of the TopoStats functions and libraries, the source code, installation instructions and a tutorial dataset are all freely available at: https://github.com/afmstats/TopoStats.

#### 2.1.2 Overview of TopoStats program

TopoStats takes raw AFM data as input, performs basic editing of the images to remove typical imaging artefacts (Figure 1A, B, C), and identifies individual molecules (Figure 1D) using Gwyddion functions. TopoStats then automatically generates backbone traces for each identified molecule (Figure 1E) and computes the contour length of circular or linear molecules without any user input. TopoStats generates length distributions for all identified molecules, and outputs this information as text files (.json files) and plots (Figure 1F) which we have used to analyse conformation of a range of biomolecules. Using our setup, TopoStats automated processing is reasonably fast: for a typical 512×512 pixel image, TopoStats corrected the artefacts and identified molecules within the image in 0.5 s and traced the identified (n = 16) molecules in 3.3 s (figure 1) on a commercially available laptop.

**Figure 1:**
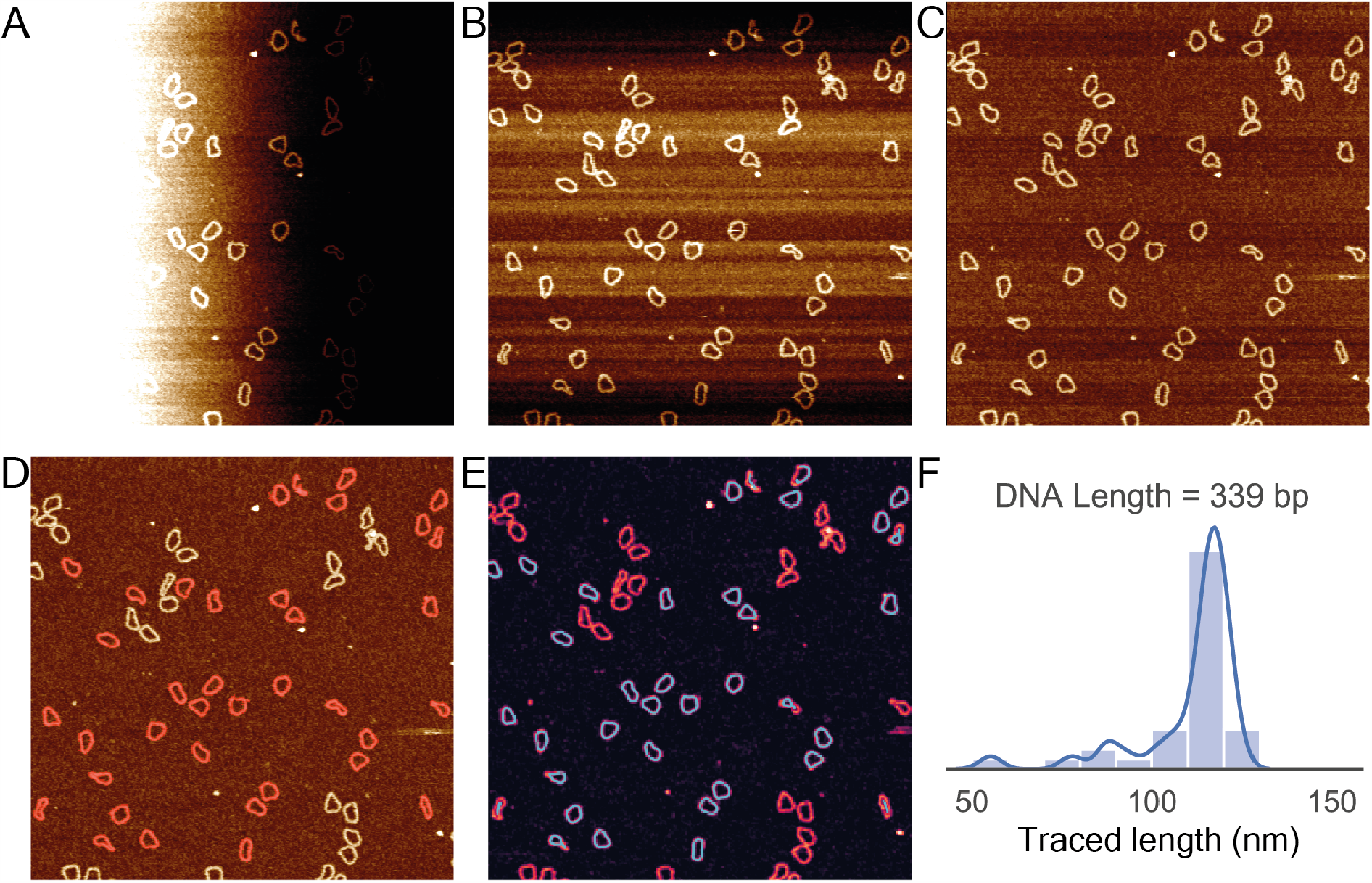
Illustration of the sequential image processing and tracing steps undertaken by TopoStats for a raw AFM image of 339 base-pair DNA minicircles. A) The original Z-scanner positional values output by the AFM, note the severe image tilt occurring due to non-perfect alignment between the sample surface and the AFM tip. (B) The tilt corrected version of the AFM image shown in (A). (C) The z-axis offset corrected version of the image shown in B. (D) The fully corrected AFM image with the identified molecules shown in red. (E) The same AFM image with overlaid molecular traces in cyan (F) A histogram of the contour lengths (nm) for each measured DNA minicircle calculated from the traces shown in E.

#### 2.1.3 AFM image correction

Distortions in raw AFM images were corrected using functions from the Gwyddion Python library ‘pygwy’. First, we used first order polynomial subtraction (i.e., plane subtraction) to remove image tilt with the Gwyddion ‘level’ function (figure 1A, B). Secondly, artefactual height (z) variations between fast scan (x-axis) line profiles were corrected by median background subtraction for each line using the Gwyddion function ‘align rows’, essentially ensuring that adjacent scan lines have matching heights (figure 1B, C). Remaining image corrections were removed using the automated Gwyddion function ‘flatten base’, which uses a combination of facet and polynomial levelling with automated masking (figure 1D). Finally, we offset the height values in the image such that the mean pixel value (corresponding to the average height value of the surface) was equal to zero. High frequency noise was removed from images using a gaussian filter (σ = 1 pixel). We found this approach sufficient for all images shown in this study, however challenging, complex or unusual samples may require additional corrections.

#### 2.1.4 Molecule Identification

TopoStats uses pygwy’s automated masking functions to identify molecules on the sample surface. In this approach, each molecule is identified using a uniquely labelled mask (grain). The positions of these grains are defined by identifying clusters of pixels by height values that deviate from the mean by a user defined value, using pygwy’s ‘datafield.mask_outliers’ function. We found a height threshold value of 0.75 − 1σ to be optimal for most samples (with 3σ corresponding to a standard gaussian). This approach initially identifies all features with heights that deviate sufficiently far from the mean surface: single molecules, clusters of molecules or aggregates and arbitrary surface contaminants. For some samples, this threshold value needs to be carefully tuned by the user, as described for a range of biomolecules in section 3.3.

To refine our grain selection to include only single molecules we employed a simple approach to remove both clusters/aggregates (large objects) and surface contaminants (typically small objects). The median area for all grains is determined and grains that have an area +/-30% of this median value are removed. An additional pruning step removes grains that contain pixels that lie on the image borders.

##### 2.1.4.1 Saving grain information

We save out the grain statistic information obtained using Gwyddion’s pygwy functions to a “.json” file, situated in the root folder and named as the root folder i.e. “myfolderofdata.json”. The grain statistic information is as follows: projected area, maximum height, mean height, minimum height, pixel area, area above half height, boundary length, minimum bounding size, maximum bounding size, centre x and y coordinates, curvature, mean radius, and ellipse angles.

#### 2.1.5 TopoStats Tracing

To implement molecule tracing in TopoStats we developed our own Python tracing library for generating smooth traces of each molecule identified as a Gwyddion grain (figure 1D, figure 2A, B). We also implemented functions in TopoStats for basic analysis of the traces (e.g. computing molecular contour length) and for visualising traces. These traces can be saved as text files, to facilitate visualisation, analysis and processing using a given user’s preferred software packages or home-written scripts.

**Figure 2:**
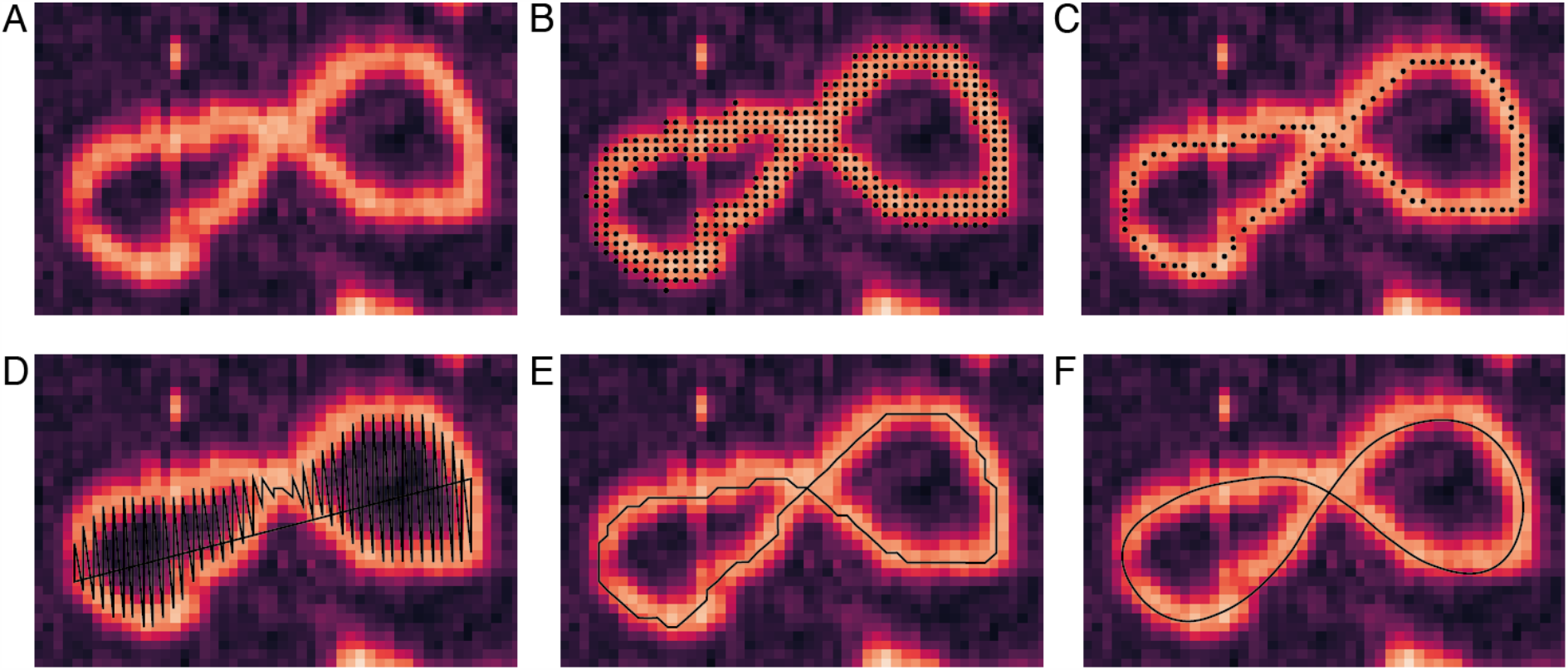
Representative image sequence showing the steps in the tracing process for an individual DNA molecule.(A) The original topographical image of a DNA minicircle. (B) The automatically generated Gwyddion grain (shown as black dots) overlaid on the DNA molecule. (C) The skeleton generated using our customised skeletonisation algorithm. Points in the skeleton are shown as black dots. (D) The cartesian coordinates for the skeleton are extracted using NumPy functions, note that the sequence of the coordinates leads to a nonsensical line trace connecting these coordinates (black line). (E) The corrected cartesian coordinates of the trace that now follows the trajectory of the underlying molecule. (F) The final smoothed trace generated by parametric splining.

TopoStats tracing is implemented using the NumPy [14] and SciPy [15] Python libraries. The tracing process is composed of 5 basic steps: firstly, the Gwyddion grain (figure 2B) is “skeletonised” into a single pixel wide binary representation of the geometric centre along the molecular backbone (figure 2C). Secondly, the positions of each pixel in the binary skeleton are extracted as cartesian coordinates (figure 2D). This initial coordinate array must be reordered such that the coordinates follow the path of the traced molecule (figure 2E). These trace coordinates are then adjusted such that they follow the highest path along the backbone of the underlying molecule. This adjusted trace is then smoothed by splining (figure 2F) to produce the final molecular trace which can be saved as a text file.

##### 2.1.5.1 Producing a rough binary skeleton

We used a modified version of the established “Zhang and Shuen” skeletonisation algorithm [16] to transform each Gwyddion grain (figure 3A) into a single pixel wide skeleton (figure 3B, C). Our adapted skeletonisation algorithm initially follows exactly the Zhang and Shuen approach: each grain is iteratively thinned by evaluating the local environment (a 3 x 3 grid) for each pixel (figure 3D), those pixels identified to be at the grain boundary are deleted whilst those at skeleton ends or required to maintain connectivity are not. We extended this process by including two additional “pruning” steps after initial skeletonisation: firstly to delete “redundant” pixels in the skeleton and secondly to remove branches that emanate from the skeleton (figure 3C). The method for identifying and removing these redundant pixels and skeleton branches is described in detail below.

**Figure 3:**
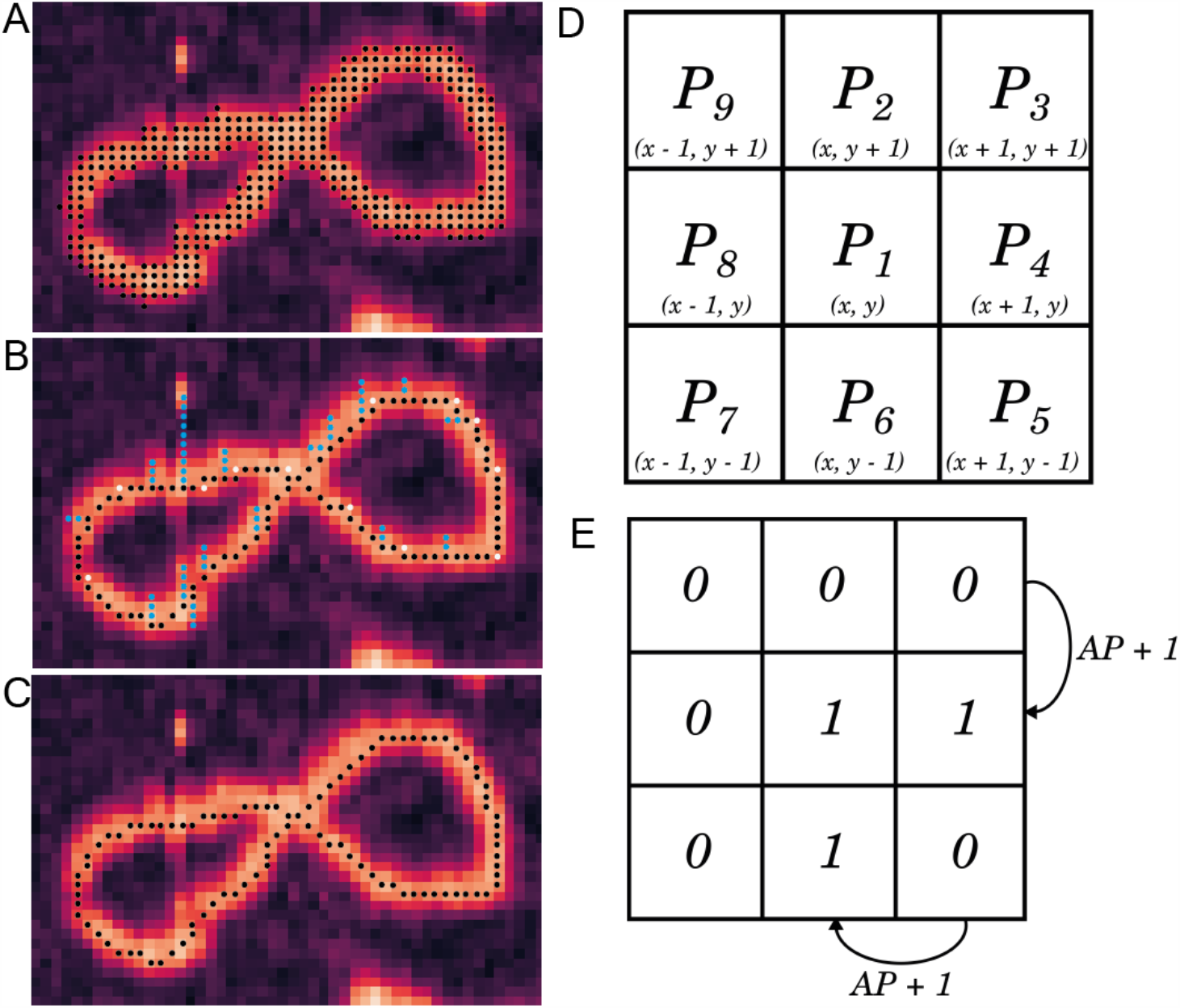
Schematic description of the skeletonisation function. (A) Example AFM image showing a DNA minicircle with the Gwyddion grain overlaid as black points. (B) A representative skeleton produced using the Zhang and Shuen approach in which branches (blue points) and redundant points (white points) can be seen within the trace. (C) The finalised skeleton with all branches and redundant points removed. (D) The naming convention for pixels within a 3×3 grid based on that used in Zhang and Shuen, 1984 as well as the reference cartesian coordinate positions for each pixel. (E) An example of a 3×3 pixel array evaluated for the (A)P1 rule.

We defined redundant pixels within the single pixel wide trace as points that were not absolutely required to maintain the connectivity and overall shape of the skeleton (figure 3B, white points), typically arising at corners in the trace. Specifically, pixels which had 2 or more direct (vertical and/or horizontal) neighbours, i.e., neighbours in P2, P4, P6 or P8 positions, were identified as redundant. Deleting these points does not break the trace connectivity, as the trace connects between the remaining points in the P2, P4, P6 or P8 positions through a diagonal connection. As this evaluation is done on the skeletonised trace, only a specific set of rules is required to identify these hanging pixels. Firstly, hanging points were identified and deleted if they satisfied the condition 1:

1. A(P1) = 2 where A(P1) is the number of [0, 1] neighbours in the (P2, P3), (P3, P4) … (P9, P2) sequence (as defined in figure 3E) and any of the following conditions 2 - 5:
2. P2 * P4 = 1
3. P4 * P6 = 1
4. P6 * P8 = 1
5. P8 * P2 = 1 Additional redundant pixels were identified and deleted if they satisfied condition 6:
6. A(P1) = 3 and any of the following conditions 7 - 10:
7. P2 * P4 * P6 = 1
8. P4 * P6 * P8 = 1
9. P6 * P8 * P2 = 1
10. P8 * P2 * P4 = 1 After redundant pixels were removed, branches from the central trace were identified and deleted (figure 3B, blue points). The Zhang Shuen skeletonisation algorithm is known to produce anomalous skeleton branches and we thus judged any short branches from the central body of the skeleton to be artifactual and removed them. We identified potential branches by locating pixels with only one neighbour within a 3×3 local environment, i.e. any pixel that satisfied condition 11:
11. B(P1) = 1 where B(P1) is the sum of all pixel values within the local 3×3 pixel environment (figure 3D). These coordinates are used to define the start of potential branches from which neighbouring pixels are sequentially added to the potential branch if they satisfy condition 12:
12. B(P1) = 2 Potential branches were deleted from the skeleton if a pixel was encountered along the potential branch that satisfied condition 13, i.e. if these branches were found to rejoin the main body of the trace:
13. B(P1) > 2

If pixels were found in potential branches that satisfied condition 11 these potential branches were judged to be linear molecules and were not deleted. This branch searching function is iterated until no branches are identified or deleted.

##### 2.1.5.2 Determination of linear and circular molecules

We used a simple approach to determine if traces were of open-ended (“linear” in DNA terminology) or closed (“closed” circular, in DNA terminology): the local 3×3 neighbour array (figure 3D) was evaluated (using condition 11) for each pixel and those with only a single neighbouring coordinate were recorded. For a closed circular trace, there will be zero coordinates with a single neighbour, whereas a linear trace will have 2 coordinates with a single neighbour (i.e. both ends of the trace).

##### 2.1.5.3 Producing an ordered trace

We extracted the cartesian coordinates of each molecule from binary skeletons (figure 4A) as a 2D NumPy array. In this procedure the coordinates are identified in ascending order along the *x*-axis and thus their sequence did not follow the trajectory of the underlying molecule and instead produced a nonsensical trace (figure 4B). As such, we reordered the coordinates, to obtain a valid representation of the traced molecule, by implementing a local-neighbour search algorithm. This algorithm iteratively identifies neighbouring coordinates from the list of “disordered” skeleton points, places the identified neighbour in the array of “ordered” coordinates and deletes this point from the list of disordered points. This approach maintains the direction of the traced molecule such that all coordinates from the skeleton are listed in a sequence that follows the trajectory of the traced molecule (figure 4C).

**Figure 4:**
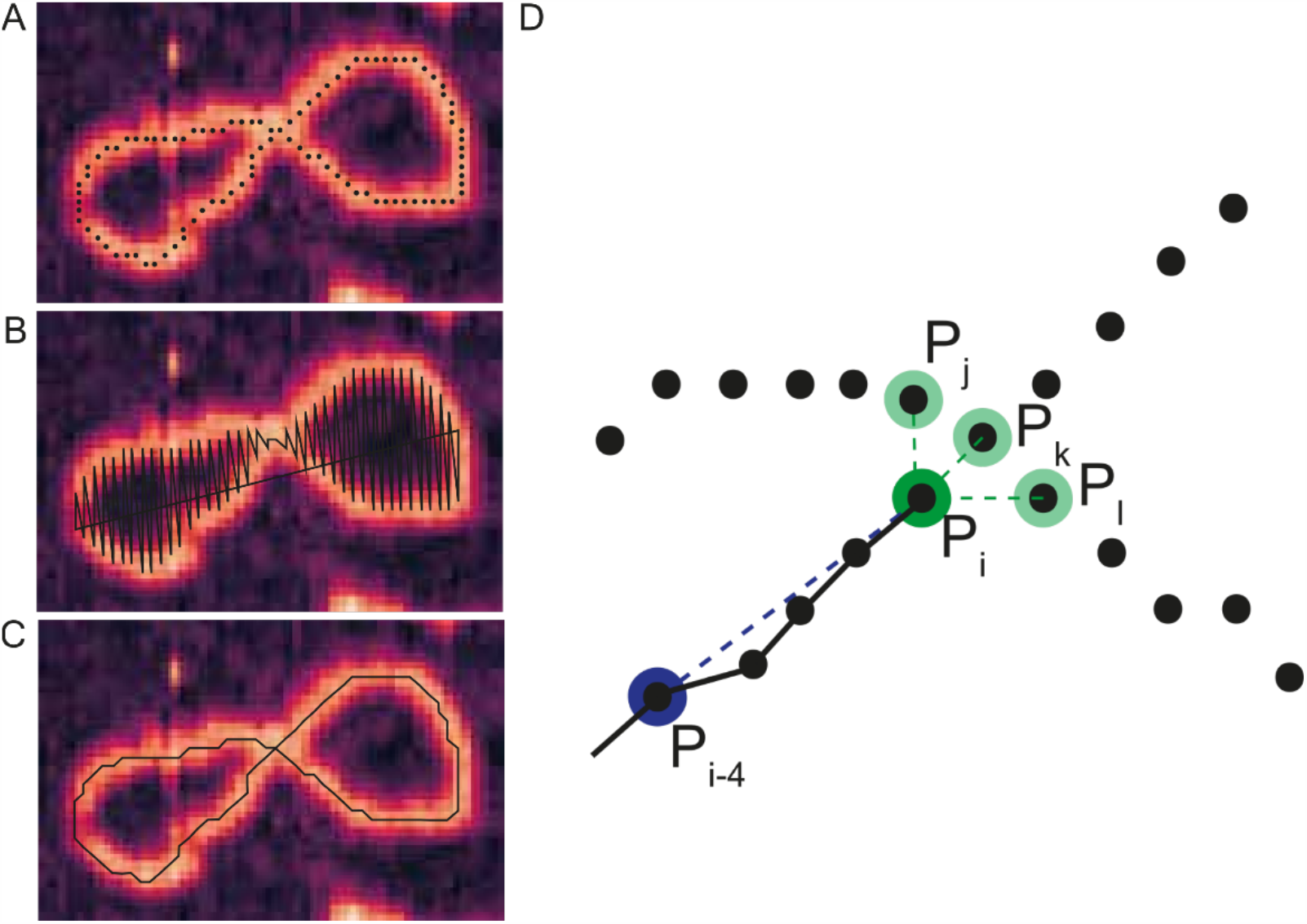
Schematic showing how the ordering process works. (A) An example image showing the pixelated binary skeleton. (B) The initial “disordered” trace in which coordinates are listed in ascending order based on the x-coordinate. Note how this trace does not follow the contours of the molecule. (C) The ordered trace that now follows the direction of the underlying molecule. (D) Diagrammatic representation of the angular search algorithm used to select the next point in the trace when multiple candidates are available. The point P_i_ is the reference point, and the reference angle is calculated using the vector between points P_i-4_ and P_i_. To distinguish between the candidate points, P_j_, P_k_ and P_l_, the angle between each candidate point and the reference point P_i-4_ is calculated. The candidate point with the vector angle most similar to that between P_i_ and P_i-4_ is accepted as the next point in the trace.

The local-neighbour search function is initiated with a sensible coordinate to start the tracing process. For linear molecules, tracing starts from one of the skeleton ends, which are identified as coordinates with only one direct neighbour (as assessed using condition 11). For circular molecules, the starting coordinate is essentially arbitrarily assigned as any of the coordinates with 2-local neighbours, ensuring that tracing does not start at a crossing of the molecule over itself. These coordinates are the first points in the “ordered” coordinate array and, crucially, are removed from the list of “disordered” skeleton points. For circular molecules, one of the 2 neighbours of the starting coordinate are arbitrarily chosen as the next point in the trace and appended to the ordered coordinate array and removed from the list of disordered points.

This starting coordinate is the first reference point (P_i_) from which the tracing algorithm identifies the next point in the trace. This next point is identified by searching the list of disordered points for neighbouring coordinates of Pi, i.e., do any coordinates lie within the 3×3 neighbourhood of Pi (figure 4D). From the first point in a linear trace, and indeed from most reference points within linear and circular traces, only one neighbouring coordinate will be present in the disordered list, which can thus be appended to the array of ordered points and removed from the disordered list. This identified coordinate then becomes the reference point for the next iteration of the tracing process. For most molecules, this simple, and fast, approach is sufficient to identify and append all points from the disordered list to the ordered array. However, a more complex method is needed to deal with reference points with multiple neighbours, which can occur when a molecule winds over itself or has a more complicated shape. At such points, the search algorithm aims to maintain the direction of the traced molecule by identifying the candidate point which deviates least from the trajectory of the coordinates in the ordered array. This is achieved by first determining the angle *θ*_i_ between the reference point P_i_ and the coordinate 3 points behind P_i_ (P_i-3_) in the ordered array. Then, the angles *θ*_i+n_ between each candidate point and the coordinate 2 points behind the reference coordinate (P_i-2_) are calculated. The candidate point whose angle *θ*_i+n_ is closest to the reference angle, *θ*_i_, is chosen as the next point in the trace, and is appended to the ordered array and removed from the disordered list.

The tracing process continues until either all the points from the disordered list are moved to the ordered array, (when the first point in the ordered array is identified as a potential next point, indicating that a circular molecule has been successfully traced), or until the reference point reaches the other skeleton end in a linear trace.

##### 2.1.5.4 Producing a fitted trace

The single pixel wide trace generated by skeletonization approximates the geometric center of the molecular backbone generated from a binary “mask” of the underlying molecule. As such, the topology of the imaged molecule has little influence on the skeleton position which can thus be an inaccurate representation of the traced molecule, particularly at sharp turns or kinks. We addressed this problem by implementing a function to adjust the trace coordinates such that they traverse a path along the highest points along the molecule (figure 5A). This function evaluates the local height profile of each trace coordinate, perpendicular to the trace direction, and adjusts the positions of each coordinate such that they lie at the highest point on the height profile (figure 5B). To avoid fitting the trace to peaks arising due to noise, the topographical image is first gaussian filtered (2 nm full-width half maxima). This improves the fit of the trace to the underlying molecule, but highly curved segments of molecules remain challenging to accurately trace.

**Figure 5:**
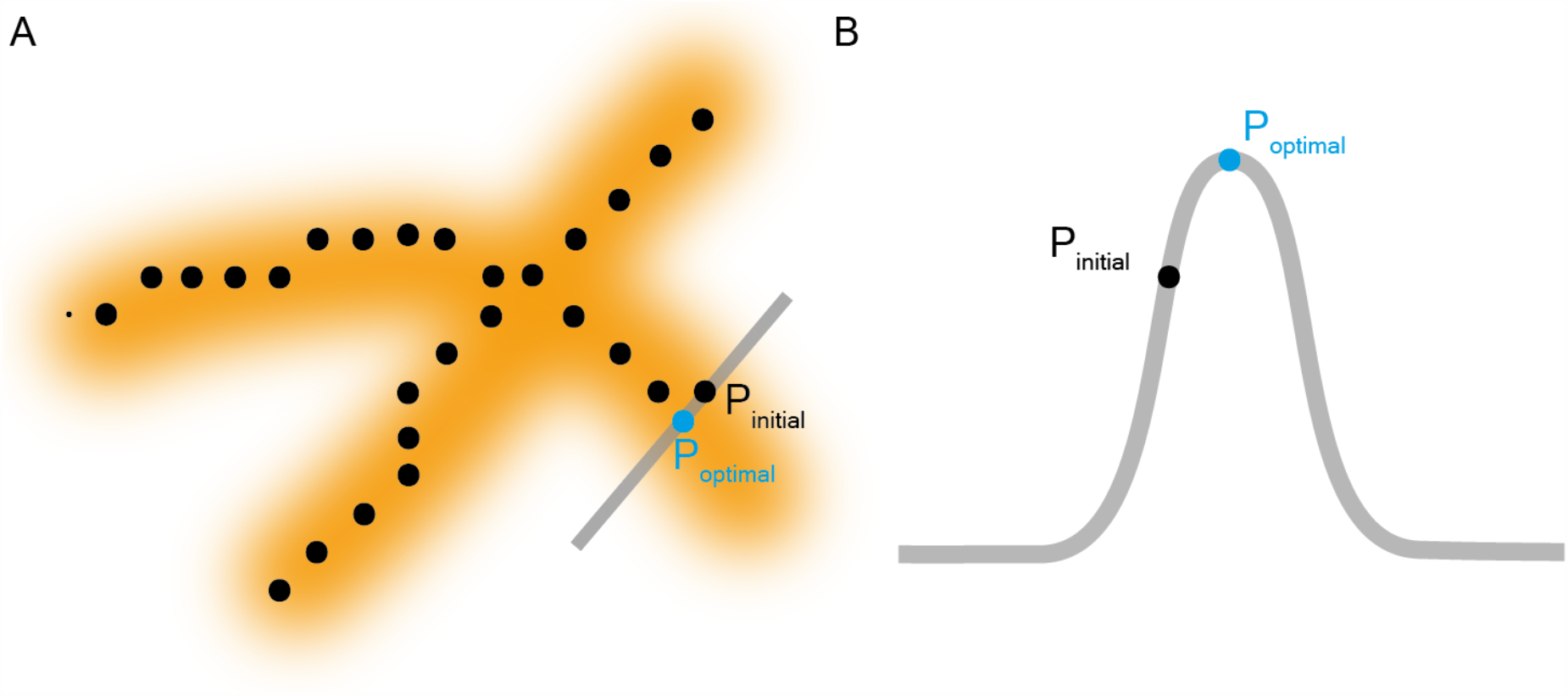
(A) Schematic of the fit-improvement protocol. The black bar represents the area that is interpolated to find the maximal height value, with the dashed red line representing the trace direction from which the perpendicular direction is determined. (B) Theoretical plot for a cross-section of height from a DNA molecule showing the original coordinate (P_intial_, black point) and the corrected coordinate (P_optimal_, blue point).

##### 2.1.5.5 Splining Coordinates

Traces generated from images with a large (>1 nm) pixel size are not sufficiently sampled to smoothly trace the underlying molecule (figure 6A). We solved this issue using parametric splining of the coordinates, to generate an interpolated trace that smoothly follows the contours of the underlying molecules. We used the SciPy interpolate functions to calculate splines. For the data presented here, the spline knots used to interpolate the traces were separated by 40Å, as an estimation of local bending. This value is defined by the user, and its value should be carefully considered based on the structural properties of the sample being investigated. To represent all points in the initial trace in the splined trace, an average of multiple independent splines is recorded (Figure 6B). For molecules with highly kinked backbones, this splining approach can underestimate local curvature. This effect can be compensated by reducing the spline knot distance or analysing the non-splined traces. It is important to note that non-splined and splined traces will have distinct contour lengths and as such may not be directly compared.

**Figure 6:**
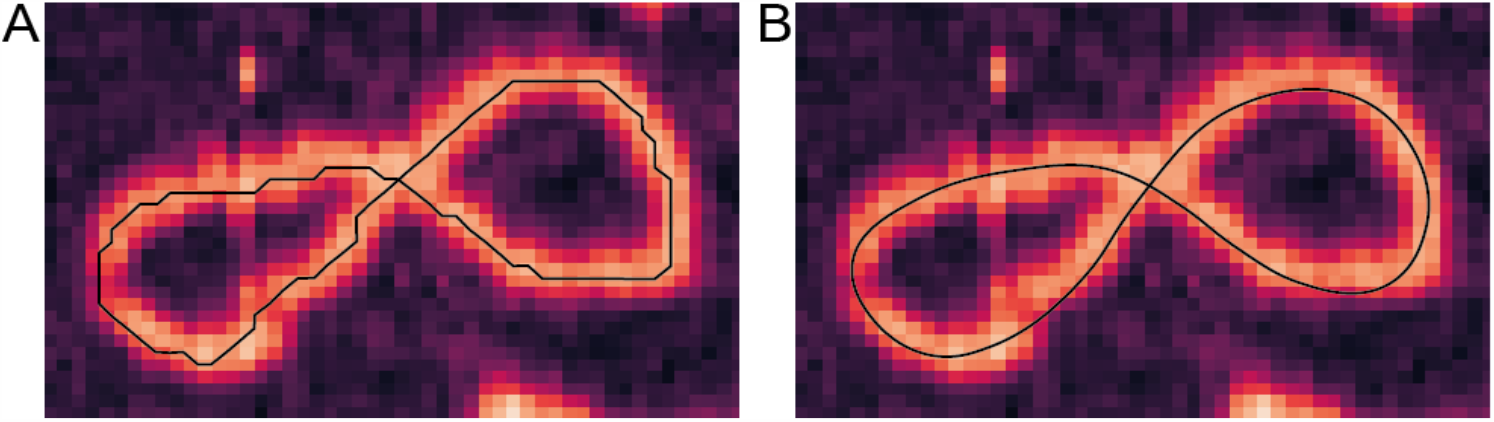
Splining smoothes out the binary traces producing a more accurate trace. (A) The original poorly sampled trace, note its coarse sampling. (B) The splined trace which smoothly follows the contours of the underlying molecule.

##### 2.1.5.6 Calculating contour length

The contour length for each trace is calculated as the sum of the vectors between all neighbouring points in the splined trace, using the following equation:

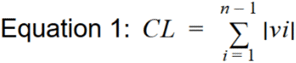

where *n* equals the number of points in the splined trace and *v*_*i*_ equals the vector between cartesian coordinates *i* and *i*+1.

##### 2.1.5.7 Saving trace information

The calculated contour length, conformation, molecule number for each traced minicircle is saved using the pandas library filename using the tracestats object to a “.json” file.

#### 2.1.6 TopoStats Plotting

TopoStats contains a ‘traceplotting’ script which uses the seaborn and matplotlib python modules [17] for data plotting. This script uses the “tracestats.json” output from the tracing as input. Data is grouped based on directory basename and is plotted as histograms, kernel density estimate (KDE) plots, and combinations of the two, in addition to violin plots.

### 2.2 Acquiring AFM images to be evaluated by TopoStats

TopoStats was designed as an effective tool for analysing molecular conformations within AFM images. It is however most effective when best practices are followed, which are explained in detail elsewhere [18]. The preparation of MAC, NuPOD and NPC samples are described in detail respectively in the literature [19][20][21][22]. As the accuracy of TopoStats is affected by the resolution of AFM imaging, we recommend following best practices for AFM imaging of soft biomaterials in solution using PeakForce Tapping mode [23][18], although sample preparation and imaging parameters may require optimisation for different samples.

#### 2.2.1 AFM Imaging

All AFM measurements were performed in liquid in PeakForce Tapping imaging on a FastScan Bio AFM system using FastScan D cantilevers (Bruker). Imaging was carried out with a PeakForce Tapping amplitude of 10 nm, at a PeakForce frequency of 8 kHz, at PeakForce setpoints of 5-20 mV, (peak forces of <100 pN). Images were recorded at 512 × 512 pixels to ensure resolution ≥ 1 nm/pixel at line rates of 3.5 Hz.

#### 2.2.2 Sample Preparation

DNA minicircles (sequences described in Appendix A) were adsorbed onto freshly cleaved mica specimen disks (diameter 6 mm, Agar Scientific, UK) at room temperature, using Ni^2+^ divalent cations. 20 μL of 3 mM NiCl_2_, 20 mM HEPES, pH 7.4 buffer solution was added to a freshly cleaved mica disk. 5-10 ng of DNA minicircles were added to the solution and adsorbed for 30 minutes. To remove any unbound DNA, the sample was washed four times using the same buffer solution.

## 3. Results and Discussion

We designed TopoStats for fast and automated structure analysis of biomolecules from AFM images. Key to this is accurate backbone tracing of polymers and oligomers, and subsequent contour length measurement and conformation determination. We used four conditions to evaluate TopoStats function, each of which we deemed essential for its widespread use. Firstly, we aimed to successfully identify the vast majority (∼90%) of available molecules that appeared isolated in the AFM images, including those from suboptimal images containing surface contaminants and aggregates. Secondly, we aimed to produce accurate traces. Thirdly, we aimed to distinguish between distinct conformations within a mixed population and, finally, we aimed to have TopoStats be versatile enough to identify and trace a range of biomolecules, without extensive optimisation and specialisation for distinct samples.

### 3.1 TopoStats for image processing and contour length determination

A key functionality of TopoStats is accurate identification and tracing of molecules from suboptimal images (those containing aggregates or surface contaminants). This facilitates faster data processing for the user as reliable molecule identification and tracing, including from poor images, reduces the need for manual inspection of each processed image. Additionally, optimising a sample to perfect homogeneity is not trivial and is often time consuming, and for some samples is not possible (Figure 7). Being able to extract useful information from suboptimal images thus facilitates AFM studies of more complex (and potentially interesting) samples and could save valuable lab time spent on sample optimisation. Here, we use two DNA minicircle samples (256 bp and 339 bp in length) to demonstrate that TopoStats can successfully identify and trace molecules from “ideal” images (339 bp sample) and from poorer images, containing aggregates and small surface contaminants (251 bp sample). To check the completeness of molecule identification in TopoStats, we also manually counted the number of isolated, non-touching DNA molecules in the images to compare to the number identified by TopoStats.

**Figure 7:**
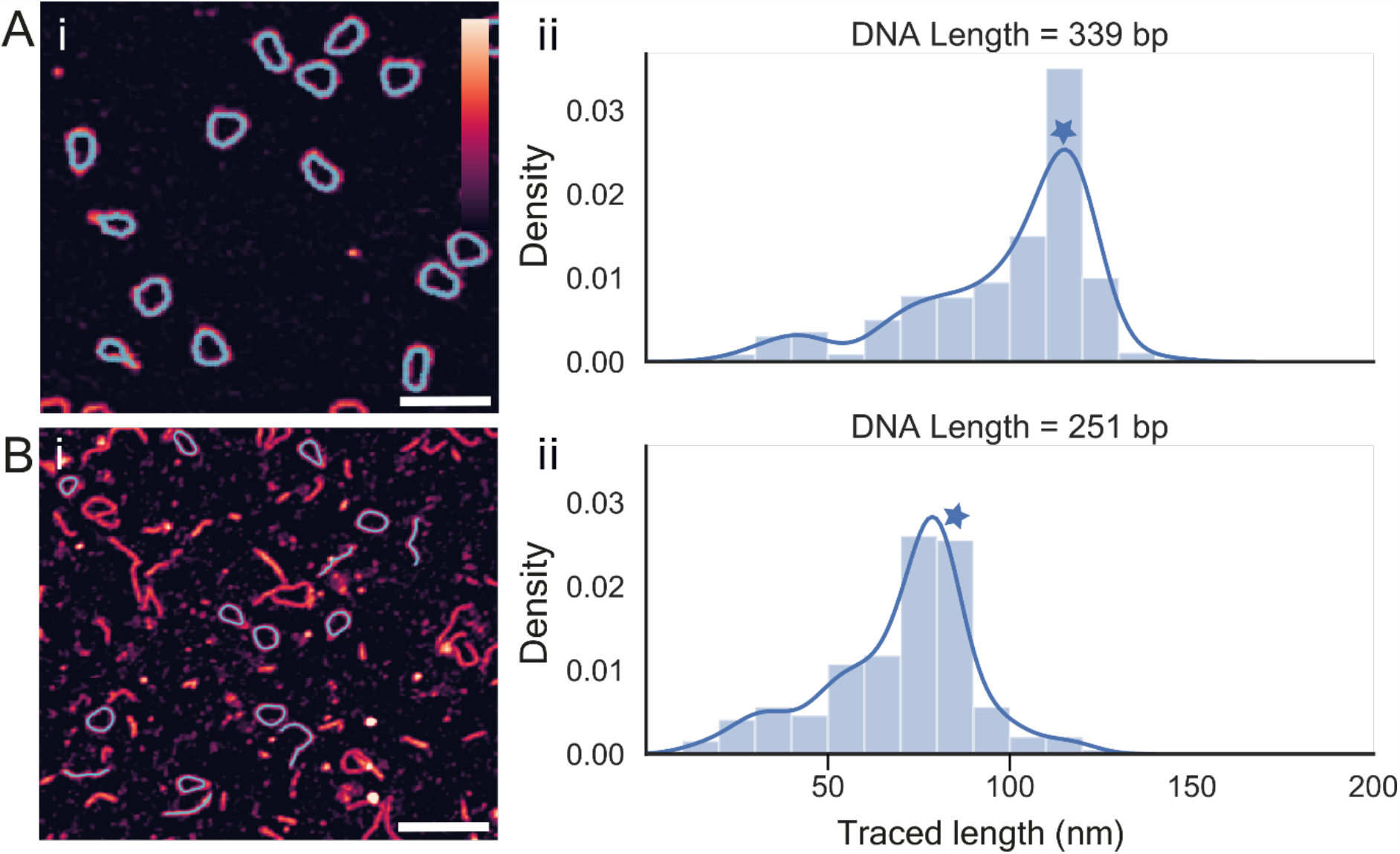
TopoStats tracing of a mixed set of images. For each dataset an (i) example AFM image is shown DNA traces overlaid in cyan and (ii) a histogram of the contour lengths. (A) 339 bp minicircles. (B) 251 bp minicircles. Blue stars represent the predicted contour lengths for each sample. Scale bars: 100 nm, vertical colour scale (inset colour bar in A): 3 nm.

Circular 339 bp DNA molecules were prepared by collaborators (appendix 1), immobilised on a mica surface, imaged with the AFM and the output raw data was analysed by TopoStats. Processed images showed a very clean sample, with essentially no aggregates or surface contaminants (figure 7Ai) facilitating excellent molecule identification: 99% of all single molecules were identified (415 of 419 molecules) and traced (figure 7Ai). The contour length histogram for 339 bp minicircles showed a well defined peak centred on the expected contour length of 115 nm (figure 7Aii). Despite the abundant presence of significant surface contaminants in the 251 bp sample, evident in the images (Figure 7Bi), this dynamic of successful molecule identification and tracing was repeated. Given the difficulty in visually distinguishing between small DNA fragments and linear DNA molecules in this sample, we only counted and compared the number of circular molecules, to minimise human bias. By this metric, TopoStats successfully identified and traced 84% of all visible molecules (figure 7Bi): 111 of 132 complete molecules. Plotting the measured contour lengths measured from these traces as a histogram showed virtually all traced molecules were full DNA minicircles: the histogram has a well-defined peak at the position of the expected contour length (85 nm for a 210 bp molecule), whilst there are comparatively few traces with shorter contour lengths (Figure 7Bii).

Given this apparent accuracy in contour length measurement for 339 and 251 bp minicircles, we further explored TopoStats tracing and contour length measurement using an expanded range of DNA minicircles samples: specifically, 116, 194, 251, 339, 357 and 398 bp. These DNA minicircles are ideal for testing TopoStats tracing accuracy as their tunable length (defined by the number of base pairs) gives a theoretical contour length (0.34 nm/bp), which can be compared to the measured contour length produced by TopoStats. The 116, 194, and 357 bp minicircles were prepared by annealing oligomers of ssDNA whilst the 251, 339 and 398 bp minicircles were prepared in bacteria by *λ*-integrase recombination (251, 339) and xer recombination (398). The 398 bp minicircles are natively negatively supercoiled, all other species are relaxed or nicked.

The DNA minicircles were prepared by collaborators (appendix 1) and immobilised on mica (as described in section 2.2.2), imaged with the AFM and the output raw AFM data was analysed with TopoStats. Examining images from each sample with overlaid traces showed that TopoStats was able to generate good traces for each construct, using default parameters. These traces followed the distinct geometries of each sample, arising from their specific lengths and production methods. For example, the shorter DNA minicircles are highly constrained by their length, which is close to the DNA persistence length (50 nm) for 194 bp (66 nm theoretical length) minicircles, and below the persistence length for 116 bp (39 nm theoretical length) minicircles. These samples were visualised as tightly compact circular conformations (Figure 8Ai-ii). This contrasts with the longer DNA minicircles (339 bp and above), which are not restricted by the persistence length and can form more complex conformations with fluctuating local curvature, whose contours are followed by the TopoStats traces (Figure 8Aiv-vi).

**Figure 8:**
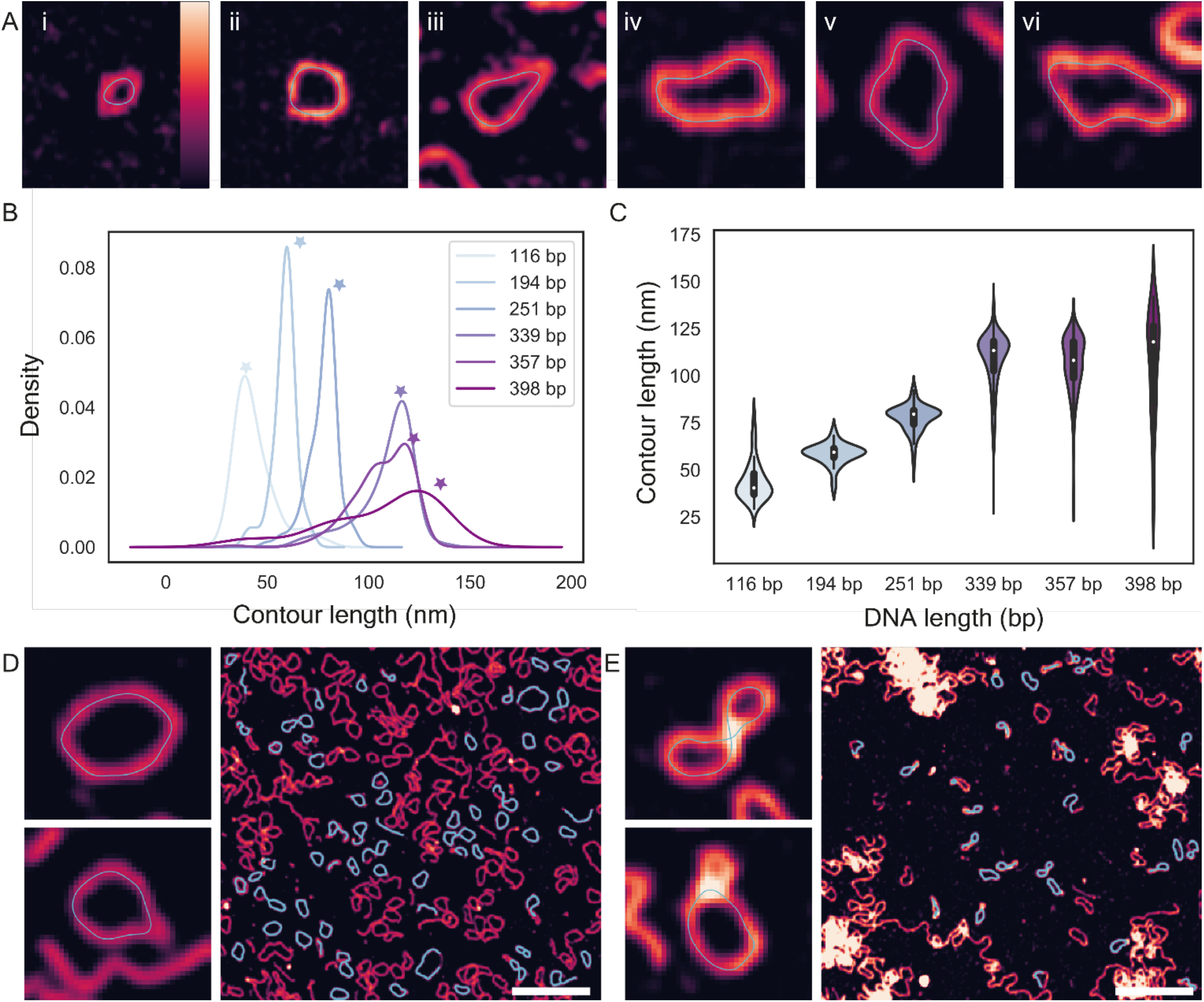
(A) Example traces (blue lines) for DNA minicircles of each length (left to right): 116 bp (i), 194 bp (ii), 256 bp (iii), 339 bp (iv), 357 bp (v), 398 bp (vi). Image widths: 80 nm, all images. Vertical scale: 6 nm (all images).(B) Kernel Density Estimate (KDE) plot showing the distributions for the measured contour lengths for each separate DNA minicircle population. Stars indicate the expected contour lengths for each sample. (C) Violin plot showing the distributions for the measured contour lengths for each separate DNA minicircle population. The median measured contour lengths are shown as white points, and correspond to 40, 59, 80, 113, 108, 118 nm, respectively. (D) Traced images from the 357 bp DNA minicircle population, note the distinct sizes of the minicircles in the top and bottom insets. (E) Traced images from the 398 bp DNA minicircle population. Scale bars: 200 nm, Vertical colour scale (inset colour bar in A): 3 nm. Images of individual DNA minicircles are 80 nm wide.

We used these TopoStats traces to calculate contour lengths for each molecule and visualised the distributions from each construct as a KDE plot (figure 8B). This distribution shows clear peaks for each species whose position increases in line with the increasing length of the DNA minicircles, and thus the theoretical contour length. We then used violin plots to better visualise the measured contour length distributions within each minicircle population (Figure 8C). These plots showed broader contour length distributions for longer constructs (339, 357 and 398 bp samples) compared to the shorter minicircles (116, 194 and 251bp) with the 357 and 398 bp samples having particularly broad distributions. The 357 bp distribution appears bimodal, with the main peak centred at ∼120 nm with a second population at ∼100 nm (figure 8A, B). We hypothesise that this minor peak is caused by an artefact in the tracing process, which produced shorter traces around highly kinked sites (Figure 8D). At highly kinked sites, the skeletonization algorithm produces a branch-like linear trace emanating from the main body, resulting in a tennis racket shaped trace. This linear branch is removed by TopoStats after skeletonization (as described in section 2.1.5.5), as it is not representative of the underlying structure. A similar broader distribution of contour lengths arises from tracing errors in the 398 bp sample. Examining the traces revealed that these errors are caused by the complexity of the minicircle conformations: the longer 398 minicircles are natively negatively supercoiled, which can lead to more compact structures that writhe (fold over on themselves) [6]. These conformations are inherently more difficult to trace, as the path of the DNA polymer is much less clear, leading to some incorrect or incomplete traces (Figure 8B), which causes a broadening of the contour length distribution. Reliably tracing these writhed (crossed) and more complex minicircle conformations should be feasible within our TopoStats framework but will likely require additional functions within the tracing modules that are specialised to deal with these complicated shapes. This is an area of current development.

With the trend established between contour length distribution and minicircle base pair length (Figure 8A, B), we next calculated the “average” (peak) measured contour length for each sample. We used the maxima of the probability distribution for each species to calculate this “average” value, as shorter DNA fragments bias the mean and median value. The measured contour lengths are listed in table 1, alongside the expected contour length (calculated based on the length in bp) and number of identified molecules. For all minicircles, there was good agreement between the peak measured contour length and the theoretical contour length: the expected length was within the noise range of the measured average for each sample. Indeed, the peak measured contour length deviated by a maximum of 6 nm from the expected value for all samples, excluding the 398 bp minicircles whose tracing was inhibited by their complex shape.

**Table 1:** Lengths of traced molecules. Expected contour lengths were calculated by multiplying the DNA length in base pairs by 0.34 nm (the helical rise of one DNA base-pair). Errors quoted are standard deviation.

| DNA length (bp) | Expected contour length (nm) | Peak measured contour length (nm) | Number of identified molecules |
| --- | --- | --- | --- |
| 116 | 39.4 | $39 \pm 10$ | 48 |
| 194 | 66.0 | $60 \pm 6$ | 51 |
| 251 | 85.3 | $80 \pm 7$ | 111 |
| 339 | 115.3 | $116 \pm 15$ | 415 |
| 357 | 121.4 | $118 \pm 14$ | 166 |
| 398 | 135.3 | $124 \pm 28$ | 36 |

Overall, this analysis demonstrates TopoStats’ capability for fully automated image correction, molecule identification and tracing from AFM images, of varying quality. For each sample, a high proportion of all molecules were identified and successfully traced (>85% of all isolated single molecules). These traces were generally very accurate, as shown by the similarity between the peak measured contour length and expected contour length, defined by the length of the minicircles in base pairs. The exception to this was the natively negatively supercoiled 398 bp sample, whose more complex shape did prove challenging for TopoStats tracing.

### 3.2 TopoStats automated determination of conformational state

Having established that TopoStats accurately measured DNA minicircle contour lengths, we next showed that TopoStats could accurately identify distinct conformations (linear and circular) within a mixed population. To do this, we used TopoStats to determine the success of a DNA annealing reaction for 194 bp minicircle construct. AFM images of an annealed DNA minicircle sample were analysed with TopoStats to determine the proportion of successfully annealed (circular) DNA molecules compared to those that did not anneal (linear molecules).

Circular 194 bp DNA molecules were prepared by collaborators, immobilised on a mica surface and imaged with the AFM (Figure 9A). Using TopoStats, we identified and traced 127 DNA molecules from 19 AFM images. Of these, 41% of DNA molecules were successfully annealed (circular) whilst 59% remained linear. Manual inspection of these images revealed a further 6 DNA molecules that had not been identified by TopoStats, 4 circular and 2 linear molecules. To further explore the differences between the linear and circular molecules within the sample, we calculated the contour lengths for each circular and linear molecules and plotted their respective distributions independently as a violin plot (Figure 9B). This showed a markedly broader contour length distribution for the linear molecules compared with the annealed circular molecules. This was reflected in the standard deviation around the mean contour lengths. Here, we used the mean contour length as we did not observe bimodal distribution for either population. The mean contour lengths and standard deviations were 55 ± 14 nm (N = 51) for linear molecules and 58 ± 6 nm (N = 76) for circular molecules. The average contour length for all traced molecules was 56 ± 12 nm. The distribution in the circular sample is narrower compared with the linear molecules as only correctly annealed and assembled molecules can form the closed circular conformations. In contrast, the linear population includes all fragmented and incorrectly annealed molecules, or those degraded by some means. It is also possible that some of this broader distribution arises due to tracing errors, like those described above (Figure 8D). Through this simple example, we show the accuracy of molecular conformation identification in TopoStats and its potential for more detailed analysis of the separated populations. We envisage this capability to be useful for more complex analysis, for example in exploring and visualising the activity of DNA nicking enzymes.

**Figure 9:**
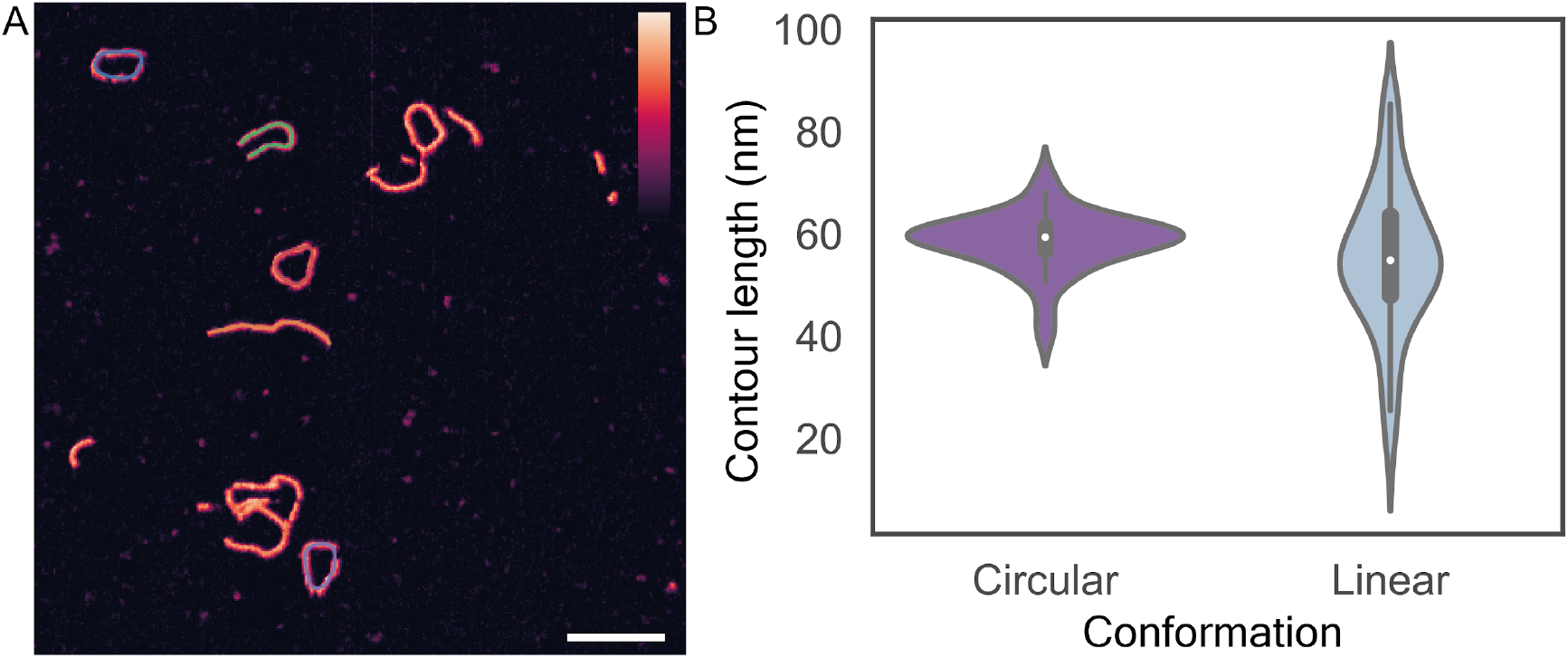
AFM analysis of DNA minicircle conformation, identifying and tracing both linear and circular molecules automatically. A) AFM image of DNA minicircles, with individual molecules traced by TopoStats. Scale bar: 50 nm, vertical colour scale (inset colour bar in A): 3 nm. B) Violin plot showing the contour length distribution for both circular (length 58 ± 6 nm, N = 51) and linear (length 55 ± 14 nm, N = 76) molecules.

### 3.3 Assessing TopoStats tracing of other Biological Molecules

Having demonstrated TopoStats’ effectiveness for identifying, tracing and reporting on the conformation of individual DNA molecules, we next explored its versatility, by tracing three distinct molecular assemblies. These were: the membrane attack complex (MAC), a hetero-oligomeric pore forming protein complex that forms circular pore assemblies in bacterial membranes. A DNA-origami biomimetic ring, NuPOD (NucleoPorins Organised on DNA), which was designed as a small synthetic mimic of the nuclear pore complex (NPC) as well as the NPC itself, a massive ring-like protein complex embedded in the nuclear membrane. These three assemblies encompass native purified protein assemblies (MAC), synthetic DNA assemblies (NuPOD) and native biological membranes extracted from cells (NPC embedded in nuclear envelope). We applied TopoStats to automatically identify individual MAC, NuPOD and NPC complexes from representative images, to assess its usefulness for these samples. For each sample, the only TopoStats parameter that needed to be optimised was the height threshold used to identify particles (section 2.1.4), as well as the size of the box used to crop individual molecules (Figure 10 A, B, C respectively).

**Figure 10:**
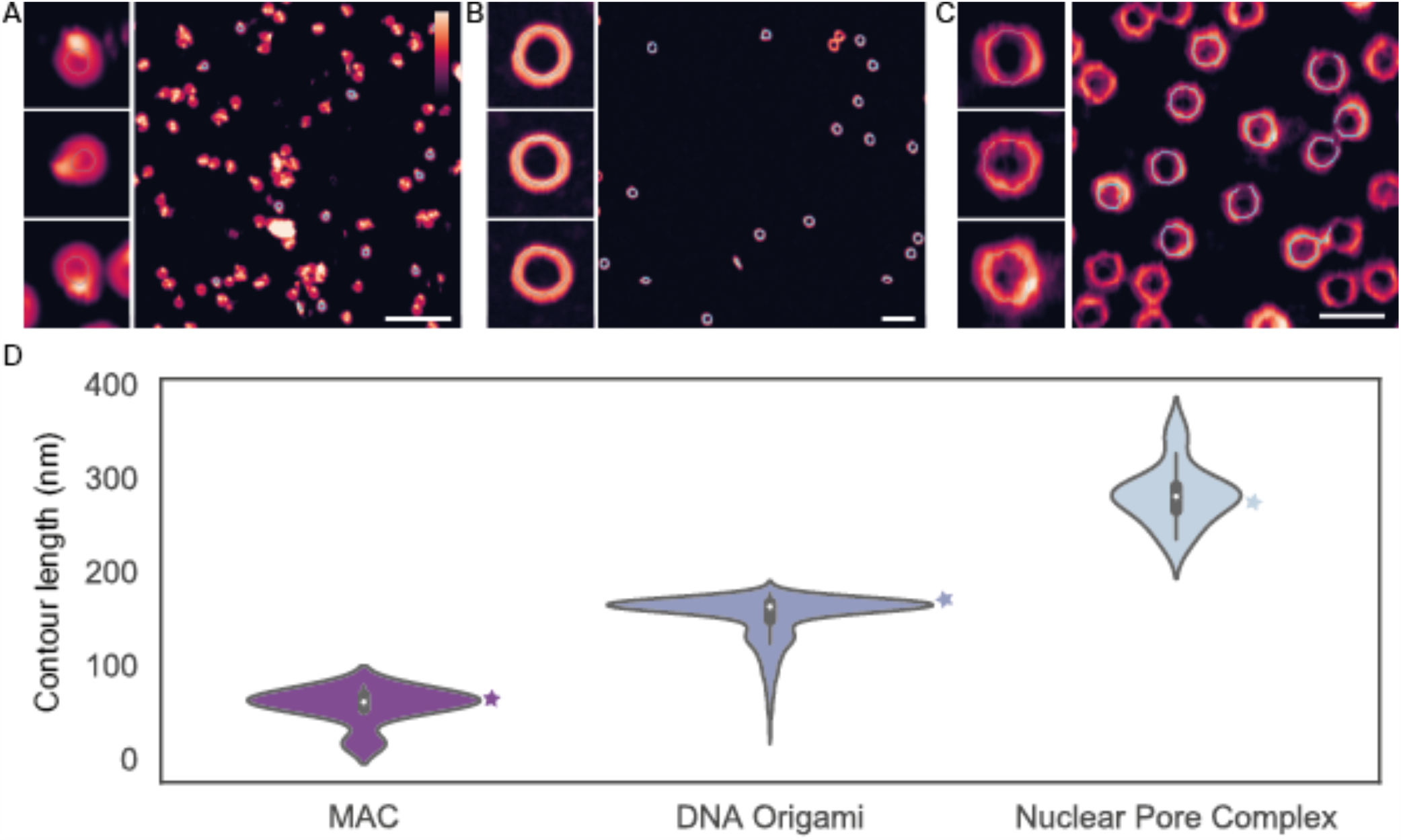
TopoStats automated tracing of A) the membrane attack complex (MAC) protein pore, B) NuPOD DNA origami rings and C) the nuclear pore complex (NPC) (C). D) Traced lengths were plotted for both assemblies with contour lengths were determined as for the 60 ± 8 nm for the MAC and 166 ± 9 nm for DNA origami determined and 287 ± 21 nm for the NPC (N = 13, 456 and 15) respectively. Stars indicate the expected contour length. Scale bars are 200 nm, cropped images are 80 nm (A), 120 nm (B) and 200 nm (C) wide. Vertical colour scale (inset colour bar in A): 20 nm (A, B) 50 nm (C). Errors quoted are standard deviation.

As with DNA minicircles, TopoStats showed excellent identification rates for the NuPOD sample, in which 97% (858 of 879 identified) of all molecules were identified, and the NPC, in which 96% were identified (24 of 25). The identification rate was poorer for the MAC where just 68% of MAC pores (13 of 19) were identified. This could be attributed to the higher height thresholding required to facilitate successful tracing of the MAC pore, and the fact that these molecular assemblies are prone to clustering. As the MAC has a very small lumen, if the entire pore is selected using a lower height threshold, the ring appears as a circle without a lumen. These circular masks are skeletonized into a single point. These single point traces are identified and discarded by TopoStats. The measured contour lengths of the assemblies were: 60 ± 8 nm for the MAC, 158 ± 8 nm for the NuPODs, and 287 ± 21 nm for the NPC (N = 13, 858 and 24 respectively). We calculated theoretical contour lengths for each sample using known pore diameters from previous studies [21][22][24], which were 63 nm (MAC), 170 nm (NuPOD) and 267 nm (NPC). In each case, the measured contour length from TopoStats showed good agreement to those from literature, demonstrating TopoStats is a versatile tool capable of producing accurate traces from a range of samples and substrates.

## 4. Conclusions

In this study, we have demonstrated the power of TopoStats, our software package for automated AFM image correction, molecule identification and tracing. Using simple examples, such as DNA minicircles at a range of lengths, we have shown that TopoStats can identify and trace isolated molecules, providing precise measures of contour length. We have also shown that TopoStats can distinguish distinct molecular conformations (circular and linear) within a mixed population. Finally, we have demonstrated that TopoStats can be applied to a range of biomolecular assemblies, including pore-forming proteins, DNA origami rings, and large protein complexes embedded in native cellular membrane, with minimal parameter changes between these different samples.

TopoStats can be used as a platform to allow processing and analysis of AFM images across a range of samples, and environments. We expect TopoStats to be a useful tool for accelerating and simplifying image processing for many working in biological AFM. As an open-source package, we hope it will be useful as a platform to facilitate the building of more complex image processing or identification routines. TopoStats has been integrated into the “AFM-SPM” community hub for AFM and SPM software development (https://github.com/AFM-SPM/home), and we actively encourage community discussion, participation, and development. We are actively developing TopoStats’ capabilities, with a focus on integrating automated polymer statistics calculation (e.g., persistence length), and to expand tracing to more complex samples e.g., DNA-protein complexes.

## Acknowledgements

The authors would like to thank Matt Newton, James Provan, Dagmar Klostermeier, Jana Hirsch, Anthony Maxwell, Lesley Mitchenhall, Jonathan Fogg and Lynn Zechiedrich for provision of DNA minicircle samples; Bernice Akpinar, Edward Parsons and George Stanley for the provision of images of DNA origami, the Membrane Attack Complex and the Nuclear Pore complex respectively; for advice and input during the development of this software; Christopher Soelistyo for assistance with developing the tracing algorithms; and Robert Turner and the Sheffield RSE team for assistance with code development. The authors acknowledge funding from the Wellcome Trust (WT 209250/Z/17/Z) and Medical Research Council (MR/R024871/1) including a UKRI/MRC Rutherford Innovation Fellowship to ALBP.

## Author Contributions

J.G.B., A.P.J., M.T. and A.L.B.P. conceived and wrote the software with assistance from L.H., R.G. and B.W.H.; R.M. and A.L.B.P. conducted AFM experiments; J.G.B. and A.L.B.P. analysed data and wrote the paper with input from R.M., A.P.J. and M.T.; all authors read and commented on paper drafts and the final version.

## Appendix A

### DNA minicircle sequences

The DNA minicircles used in this study were prepared by collaborators, and stored in buffer solution or water at 4°C or −20 °C, at concentrations of 1-100 ng/µL prior to use. DN minicircle samples are identified by their length in base pairs (bp).

**116 bp**: DNA minicircles were prepared as detailed in [25] by self-assembly of short oligos to form a non-ligated circle of 116 bp dsDNA with a 21 bp ssDNA quadruplex forming motif contained within.

**194 bp**: DNA minicircles were prepared by self-assembly of short oligos to form a non-ligated circle of 210 bp DNA containing 194 bp dsDNA and a 16 bp ssDNA inset [26][27]. Sequence: ACTTTTTTGTGGGTTTTTGAGGCCGCGTTCAGCCTTTTTCGCCGTTTTTTGCGAATTTTTCAGTCTTTTTTGGTCCTTTTTGCGACTTTTTTCGGCGTTTTTCTGCCTTTTTTGCGTGTTTTTGACCCTTTTTTCGCAGTTTTTGGCTCTTTTTTGCAGCTTTTTAATTAAGGAGGAGGAGGAGAAGGAGATTTTTTACGCATTTTTGTC

**251 bp**: DNA minicircles were prepared as in [28][7], using lambda-integrase recombination followed by purification. Sequence: TTTATACTAACTTGAGCGAAACGGGAAGGTAAAAAGACAACAAACTTTCTTGTATACCTTTAAGAGAGAGAGAGAGAGACGACTCCTGCGATATCGCCTCGGCTCTGTTACAGGTCACTAATACCATCTAAGTAGTTGATTCATAGTGACTGCATATGTTGTGTTTTACAGTATTATGTAGTCTGTTTTTTATGCAAAATCTAATTTAATATATTGATATTTATATCATTTTACGTTTCTCGTTCAGCTTT

**339 bp**: DNA minicircles were prepared as in [29][28], using lambda-integrase recombination followed by purification. Sequence: TTTATACTAACTTGAGCGAAACGGGAAGGGTTTTCACCGATATCACCGAAACGCGCGAGGCAGCTGTATGGCGAAATGAAAGAACAAACTTTCTTGTACGCGGTGGTGAGAGAGAGAGAGAGATACGACTACTATCAGCCGGAAGCCTATGTACCGAGTTCCGACACTTTCATTGAGAAAGATGCCTCAGCTCTGTTACAGGTCACTAATACCATCTAAGTAGTTGATTCATAGTGACTGCATATGTTGTGTTTTACAGTATTATGTAGTCTGTTTTTTATGCAAAATCTAATTTAATATATTGATATTTATATCATTTTACGTTTCTCGTTCAGCTTT

**357 bp**: DNA minicircles were prepared by self-assembly of short oligos to form a ligated circle of 357 bp dsDNA. Sequence: TGGACAGCTTATCATCGATAAGCTTGCTAGCGGGCCCTGTAGGCCCACTTAACACTACAAGACCTACGCCTCTCCATTCATCATGTAACCCACAAATCATCTAAACCGTAAGTCTAAGGGCCTCCTGAGGTTTTCTCAGGAGGCCCTAATGTATAATTATGATGGGAGCCCTTCTTCTTCTGCTCGGACTCAGGCTTATACATATTTGAATGTATTTAGAAAAATAAACAAATAGGGGTTCCGCGCACATTTCCCCGAAAAGTGCCACCTGACGTCTAAGAAACCATTATTATCATGACATTAACCTATAAAAATAGGCGTATCACGAGGCCCTTTCGTCTTCAAGAGCTCTCATGT

**398 bp**: 398bp DNA minicircles were prepared by in-vitro E. coli xer recombination using the plasmid DNA substrate pSDC153, as detailed in [30]. Sequence: GGGTACCGAGCTCGAATTGACTCTAGAGGATCCCCTGAGACAACTTGTTACAGCTCAACAGTCACACAT AGACAGCCTGAAACAGGCGATGCTGCTTATCGAATCAAAGCTGCCGACAACACGGGAGCCAGTGACGC CTCCCGTGGGGAAAAAATCATGGCAATTCTGGAAGAAATAGCGCTTTCAGCCGGCAAACCggcTGAAGC CGGATCTGCGATTCTGATAACAAACTAGCAACACCAGAACAGCCCGTTTGCGGGCAGCAAAACCCGTAC TTTTGGACGTTCCGGCGGTTTTTTGTGGCGAGTGGTGTTCGGGCGGTGCGCGCAAGATCCATTATGTTAA ACGGGCGAGTTTACATCTCAAAACCGCCCGCTTAACACCATCAGAAATCCTCA

## References

[1] A. P. Nievergelt, N. Banterle, S. H. Andany, P. Gönczy, and G. E. Fantner, “High-speed photothermal off-resonance atomic force microscopy reveals assembly routes of centriolar scaffold protein SAS-6,” Nat. Nanotechnol., p. 1, May 2018, doi: 10.1038/s41565-018-0149-4.

[2] T. Uchihashi, R. Iino, T. Ando, and H. Noji, “High-Speed Atomic Force Microscopy Reveals Rotary Catalysis of Rotorless F1-ATPase,” Science (80-.)., vol. 333, no. 6043, pp. 755–758, Aug. 2011, doi: 10.1126/science.1205510.

[3] A. Pyne, R. Thompson, C. Leung, D. Roy, and B. W. Hoogenboom, “Single-Molecule Reconstruction of Oligonucleotide Secondary Structure by Atomic Force Microscopy,” Small, vol. 10, no. 16, pp. 3257–3261, 2014, doi: 10.1002/smll.201400265.

[4] T. Uchihashi et al., “Dynamic structural states of ClpB involved in its disaggregation function,” Nat. Commun., vol. 9, no. 1, p. 2147, Dec. 2018, doi: 10.1038/s41467-018-04587-w.

[5] N. Kodera, D. Yamamoto, R. Ishikawa, and T. Ando, “Video imaging of walking myosin V by high-speed atomic force microscopy,” Nature, vol. 468, no. 7320, pp. 72–76, 2010, doi: 10.1038/nature09450.

[6] H. Yamashita, K. Voïtchovsky, T. Uchihashi, S. A. Contera, J. F. Ryan, and T. Ando, “Dynamics of bacteriorhodopsin 2D crystal observed by high-speed atomic force microscopy,” J. Struct. Biol., vol. 167, no. 2, pp. 153–158, Aug. 2009, doi: 10.1016/j.jsb.2009.04.011.

[7] A. Pyne et al., “Combining high-resolution AFM with MD simulations shows that DNA supercoiling induces kinks and defects that enhance flexibility and recognition,” bioRxiv, p. 863423, Dec. 2019, doi: 10.1101/863423.

[8] W. Kühlbrandt, “The resolution revolution,” Science, vol. 343, no. 6178. American Association for the Advancement of Science, pp. 1443–1444, Mar. 28, 2014, doi: 10.1126/science.1251652.

[9] J. Zivanov et al., “RELION-3: new tools for automated high-resolution cryo-EM structure determination,” bioRxiv, p. 421123, Sep. 2018, doi: 10.1101/421123.

[10] J. Schindelin et al., “Fiji: an open-source platform for biological-image analysis,” Nat. Methods, vol. 9, no. 7, pp. 676–682, Jul. 2012, doi: 10.1038/nmeth.2019.

[11] D. Nečas and P. Klapetek, “Gwyddion: an open-source software for SPM data analysis,” Open Phys., vol. 10, no. 1, pp. 181–188, Jan. 2012, doi: 10.2478/s11534-011-0096-2.

[12] S. Konrad et al., “High-throughput AFM analysis reveals unwrapping pathways of H3 and CENP-A nucleosomes,” bioRxiv, p. 2020.04.09.034090, Apr. 2020, doi: 10.1101/2020.04.09.034090.

[13] A. Mikhaylov, S. K. Sekatskii, and G. Dietler, “DNA trace: A comprehensive software for polymer image processing,” J. Adv. Microsc. Res., vol. 8, no. 4, pp. 241–245, Dec. 2013, doi: 10.1166/jamr.2013.1164.

[14] S. van der Walt, S. C. Colbert, and G. Varoquaux, “The NumPy Array: A Structure for Efficient Numerical Computation,” Comput. Sci. Eng., vol. 13, no. 2, pp. 22–30, Mar. 2011, doi: 10.1109/MCSE.2011.37.

[15] P. Virtanen et al., “SciPy 1.0: fundamental algorithms for scientific computing in Python,” Nat. Methods, vol. 17, no. 3, pp. 261–272, Mar. 2020, doi: 10.1038/s41592-019-0686-2.

[16] T. Y. Zhang and C. Y. Suen, “A Fast Parallel Algorithm for Thinning Digital Patterns,” Commun. ACM, vol. 27, no. 3, pp. 236–239, Mar. 1984, doi: 10.1145/357994.358023.

[17] J. D. Hunter, “Matplotlib: A 2D graphics environment,” Comput. Sci. Eng., vol. 9, no. 3, pp. 99–104, May 2007, doi: 10.1109/MCSE.2007.55.

[18] A. L. B. Pyne and B. W. Hoogenboom, “Imaging DNA structure by atomic force microscopy,” in Methods in Molecular Biology, vol. 1431, Humana Press Inc., 2016, pp. 47–60.

[19] E. S. Parsons et al., “Single-molecule kinetics of pore assembly by the membrane attack complex,” Nat. Commun., vol. 10, no. 1, pp. 1–10, Dec. 2019, doi: 10.1038/s41467-019-10058-7.

[20] P. D. E. Fisher et al., “A Programmable DNA Origami Platform for Organizing Intrinsically Disordered Nucleoporins within Nanopore Confinement,” ACS Nano, vol. 12, no. 2, pp. 1508–1518, Feb. 2018, doi: 10.1021/acsnano.7b08044.

[21] G. J. Stanley et al., “Quantification of Biomolecular Dynamics Inside Real and Synthetic Nuclear Pore Complexes Using Time-Resolved Atomic Force Microscopy,” ACS Nano, vol. 13, no. 7, pp. 7949–7956, Jul. 2019, doi: 10.1021/acsnano.9b02424.

[22] G. J. Stanley, A. Fassati, and B. W. Hoogenboom, “Atomic force microscopy reveals structural variability amongst nuclear pore complexes,” Life Sci. Alliance, vol. 1, no. 4, Aug. 2018, doi: 10.26508/lsa.201800142.

[23] H. Yamashita, N. Kodera, A. Miyagi, T. Uchihashi, D. Yamamoto, and T. Ando, “Tip-sample distance control using photothermal actuation of a small cantilever for high-speed atomic force microscopy,” Rev. Sci. Instrum., vol. 78, no. 8, p. 083702, Aug. 2007, doi: 10.1063/1.2766825.

[24] A. Menny et al., “CryoEM reveals how the complement membrane attack complex ruptures lipid bilayers,” Nat. Commun., vol. 9, no. 1, pp. 1–11, Dec. 2018, doi: 10.1038/s41467-018-07653-5.

[25] B. Klejevskaja et al., “Studies of G-quadruplexes formed within self-assembled DNA mini-circles,” Chem. Commun., vol. 52, no. 84, pp. 12454–12457, Oct. 2016, doi: 10.1039/c6cc07110d.

[26] J. Valero, N. Pal, S. Dhakal, N. G. Walter, and M. Famulok, “A bio-hybrid DNA rotor-stator nanoengine that moves along predefined tracks,” Nat. Nanotechnol., vol. 13, no. 6, pp. 496–503, Jun. 2018, doi: 10.1038/s41565-018-0109-z.

[27] A. Valero-Rello et al., “Validation and implementation of a diagnostic algorithm for DNA detection of bordetella pertussis, B. parapertussis, and B. holmesii in a Pediatric Referral Hospital in Barcelona, Spain,” J. Clin. Microbiol., vol. 57, no. 1, Jan. 2019, doi: 10.1128/JCM.01231-18.

[28] J. M. Fogg et al., “Exploring writhe in supercoiled minicircle DNA,” J. Phys. Condens. Matter, vol. 18, no. 14, Apr. 2006, doi: 10.1088/0953-8984/18/14/S01.

[29] R. N. Irobalieva et al., “Structural diversity of supercoiled DNA,” Nat. Commun., vol. 6, Oct. 2015, doi: 10.1038/ncomms9440.

[30] S. D. Colloms, J. Bath, and D. J. Sherratt, “Topological selectivity in Xer site-specific recombination,” Cell, vol. 88, no. 6, pp. 855–864, Mar. 1997, doi: 10.1016/S0092-8674(00)81931-5.

